# Relative model selection of evolutionary substitution models can be sensitive to multiple sequence alignment uncertainty

**DOI:** 10.1101/2021.08.04.455051

**Authors:** Stephanie J. Spielman, Molly L. Miraglia

## Abstract

**Background:** Multiple sequence alignments (MSAs) represent the fundamental unit of data inputted to most comparative sequence analyses. In phylogenetic analyses in particular, errors in MSA construction have the potential to induce further errors in downstream analyses such as phylogenetic reconstruction itself, ancestral state reconstruction, and divergence time estimation. In addition to providing phylogenetic methods with an MSA to analyze, researchers must also specify a suitable evolutionary model for the given analysis. Most commonly, researchers apply relative model selection to select a model from candidate set and then provide both the MSA and the selected model as input to subsequent analyses. While the influence of MSA errors has been explored for most stages of phylogenetics pipelines, the potential effects of MSA uncertainty on the relative model selection procedure itself have not been explored.

**Results:** We assessed the consistency of relative model selection when presented with multiple perturbed versions of a given MSA. We find that while relative model selection is mostly robust to MSA uncertainty, in a substantial proportion of circumstances, relative model selection identifies distinct best-fitting models from different MSAs created from the same set of sequences. We find that this issue is more pervasive for nucleotide data compared to amino-acid data. However, we also find that it is challenging to predict whether relative model selection will be robust or sensitive to uncertainty in a given MSA.

**Conclusions:** We find that that MSA uncertainty can affect virtually all steps of phylogenetic analysis pipelines to a greater extent than has previously been recognized, including relative model selection.

## Background

One of the first steps in a phylogenetic analysis, and indeed any comparative sequence analysis, is to construct a multiple sequence alignment (MSA) from homologous sequences of interest. In spite of decades of advances in MSA construction algorithms, it remains difficult to confidently generate a reliable and accurate MSA that fully considers the evolutionary process, and different software platforms and/or algorithm parameterizations are known yield different MSAs of varying quality (Wong et al. 2008; Penn et al. 2010; Thompson et al. 2011; Sela et al. 2015). A substantial body of research has shown that errors in the MSA itself can influence, with the potential to produce spurious results, downstream analyses such as phylogenetic reconstruction (Wong et al. 2008; Ashkenazy et al. 2019), phylogenetic topology tests (Levy Karin et al. 2014), ancestral state reconstruction (Ashkenazy et al. 2019), divergence time estimation (Du et al. 2019), and identification of positive selection (Markova-Raina and Petrov 2011; Jordan and Goldman 2012; Privman et al. 2012).

While the effects of MSA uncertainty in phylogenetic pipelines have been heavily studied, the MSA is not the only piece of information that is inputted to phylogenetic reconstruction and other evolutionary-informed analyses. A suitable model of sequence evolution for the data at hand must also be provided (Arenas 2015). Most models of sequence evolution consider a continuous-time reversible Markov process, and there are dozens, if not hundreds, of available parameterizations for nucleotide and amino-acid models (Yang 2014). One of the most common approaches used to identify a suitable model for phylogenetic inference is relative model selection, wherein a set of candidate models are ranked according to a given goodness-of-fit measurement, and the best-fitting model is then used in the phylogenetic reconstruction (Sullivan and Joyce 2005). Although recent studies have suggested that relative model selection may not be a critical step in phylogenetic studies (Spielman and Kosakovsky Pond 2018; Abadi et al. 2019; Spielman 2020), it remains an enduring staple of most analysis pipelines. Henceforth, we use the phrase “model selection” to refer specifically to relative model selection, unless otherwise stated.

When performing model selection, the MSA of interest is generally fixed, yet there is rarely a guarantee that the MSA has precisely identified site homology and/or insertion/deletion (indel) events. Indeed, there are many MSAs which could be in theory derived from a given set of homologous sequences, but how relative model selection would perform on a different MSA version is unclear. In other words, if there were *N* different MSAs which could be generated from the same set of orthologs, it is unknown whether relative model selection will consistently identify the same best-fitting model for all *N* MSAs. Previously, Abdo et al. (2005) showed that relative model selection is robust to the guide tree used to assess model fit, but it remains unknown to what extent uncertainty in the MSA will influence the procedure. This dearth of understanding is a notable omission from both the model selection literature and from the literature investigating effects of MSA uncertainty on phylogenetic applications.

Here, we aimed to bridge this gap through a systematic analysis of whether MSA uncertainty can affect the results of model selection. We examined whether model selection is robust to MSA uncertainty across thousands of sets of natural orthologous sequences, at both the amino-acid and nucleotide levels, by interrogating whether model selections identifies different optimal models for different versions of MSAs created from the same underlying set of orthologs. We broadly found that there is potential for model selection, in particular on nucleotide data, to identify different best-fitting evolutionary models for different MSA versions created from the same ortholog set. Moreover, we find that it is challenging to predict whether model selection will be robust or sensitive to uncertainty for a given MSA. Our results demonstrate that MSA uncertainty may be even more of a pervasive influence in phylogenetic analysis pipelines than has previously been recognized.

## Results

We obtained sets of unaligned orthologs (“datasets”; Table 1) from databases Selectome (Moretti et al. 2014), including both Euteleostomi and Drosophila sequences, and PANDIT (Whelan 2006). For each dataset, we generated 50 unique MSA variants using a `GUIDANCE2`-based approach (Sela et al. 2015; Spielman et al. 2014), with one MSA designated as the reference MSA and the remaining 49 designated as perturbed MSAs. The `GUIDANCE2` algorithm generates alternative MSAs by varying the guide tree and gap opening penalties used by progressive alignment algorithms during MSA reconstruction. For each resulting MSA, we used `ModelFinder` (Nguyen et al. 2015; Kalyaanamoorthy et al. 2017) to determine the best-fitting model with each of the three information theoretic criteria AIC, AICc, and BIC. For all steps, we analyzed both the nucleotide (NT) and amino-acid (AA) versions (“datatypes”; Table 1) of each dataset. In total, for each datatype, we processed 1000 Drosophila and Euteleostomi datasets each from Selectome and 236 datasets from PANDIT, more details for which are available in *Methods and Materials*. Associated acronyms and terms are defined in Table 1.

**Table 1:** Terminology used throughout text.

| Term | Meaning |
| --- | --- |
| **MSA** | Acronym for **m**ultiple **s**equences **a**lignment |
| **Dataset** | A set of unaligned orthologous sequences |
| **Datatype** | Either nucleotide (NT) or amino-acid (AA) |
| **Dataset source** | Either Drosophila, Euteleostomi, or PANDIT |
| **Reference MSA** | The initial MSA created from a given dataset using default MAFFT settings |
| **Perturbed MSA** | A perturbed MSA generated from a given dataset using a GUIDANCE2-based approach |
| **Variant MSA** | One of any 50 unique MSAs (either reference or variant MSA) built from a given dataset |
| **Stable dataset** | A dataset with the same selected model for all variant MSAs |
| **Unstable dataset** | A dataset with at least two distinct selected models among all variant MSAs |
| $M_0$ | The most frequently identified best-fitting model for a given dataset |
| $M_{\text{ref}}$ | The best-fitting model identified for a given dataset’s reference MSA |

### Distinct models are often associated with variant MSAs

We first investigated model selection’s consistency by tabulating how many distinct best-fitting models were identified across a given dataset’s 50 variant MSAs. If model selection were robust to MSA uncertainty, we would observe the same best-fitting model for all variant MSAs. Alternatively, if model selection were sensitive to MSA uncertainty, we would observe multiple distinct best-fitting models identified across a given dataset’s variant MSAs. We refer to datasets adhering to the former scenario (one model for all variant MSAs) as a “stable” and datasets adhering to the latter scenario as “unstable” (Table 1).

Overall, we observed distinct stability patterns across datatypes and dataset source (Figure 1). Drosophila datsets tended to be more unstable, with a pronounced instability increase for nucleotide datasets compared to amino-acid datasets. By contrast, Euteleostomi datasets tended to be more stable, with a pronounced stability increase for amino-acid datasets compared to nucleotide datasets. Finally, PANDIT amino-acid datasets tended to be more stable compared to their nucleotide counterparts. Stability was broadly consistent across the theoretic information criterion used, suggesting that all criteria are similarly influenced by MSA uncertainty.

**Figure 1:**
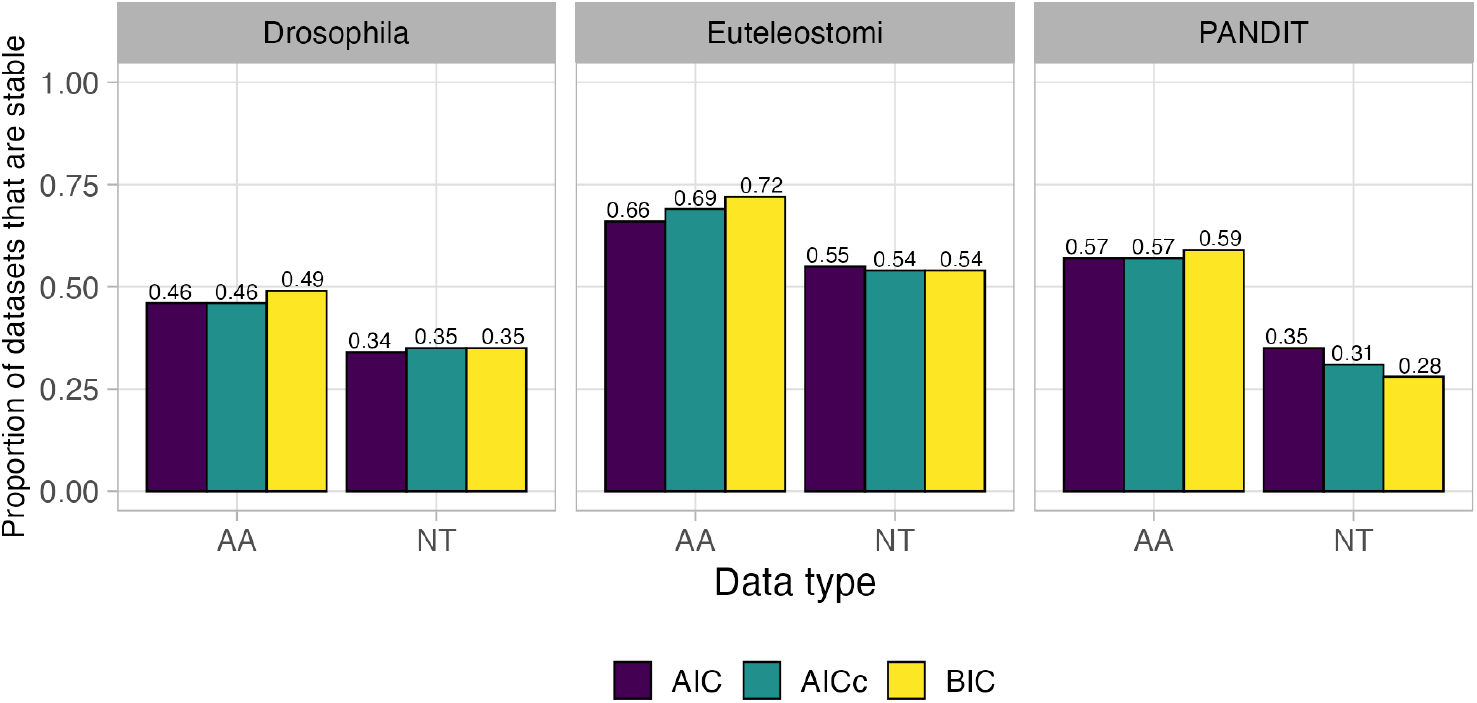
Percent of stable and unstable (defined in Table 1) datasets within each of the three data sources, Selectome Euteleostomi [1000 amino-acid (AA) and nucleotide (NT) datasets each], Selectome Drosophila (1000 AA and NT datasets each), and PANDIT (236 AA and NT datasets each). Bars are colored based on the information theoretic criterion used for model selection.

We next counted the total number of selected models identified for unstable datasets (Figure 2a for AIC results and Figure S1a,c for BIC and AICc results, respectively). Results were again broadly consistent among information theoretic criteria. Across both amino-acid and nucleotide datatypes, the vast majority of unstable datasets only had two associated best-fitting models; for example, as shown in Figure 2a, that roughly 65% of all Drosophila amino-acid datasets were associated with two distinct best-fitting models with model selection by AIC. Across all amino-acid datasets, roughly 17–34% of ortholog datasets were associated with three or more models, and across all nucleotide datasets, roughly 32–52% of ortholog datasets were associated with three or more models, demonstrating that model selection is indeed influenced by MSA uncertainty in a substantial proportion of cases. Most notably, model selection was most sensitive to MSA uncertainty on Drosophila NT datasets, of which nearly 10% of datasets selected five or more best-fitting models.

**Figure 2:**
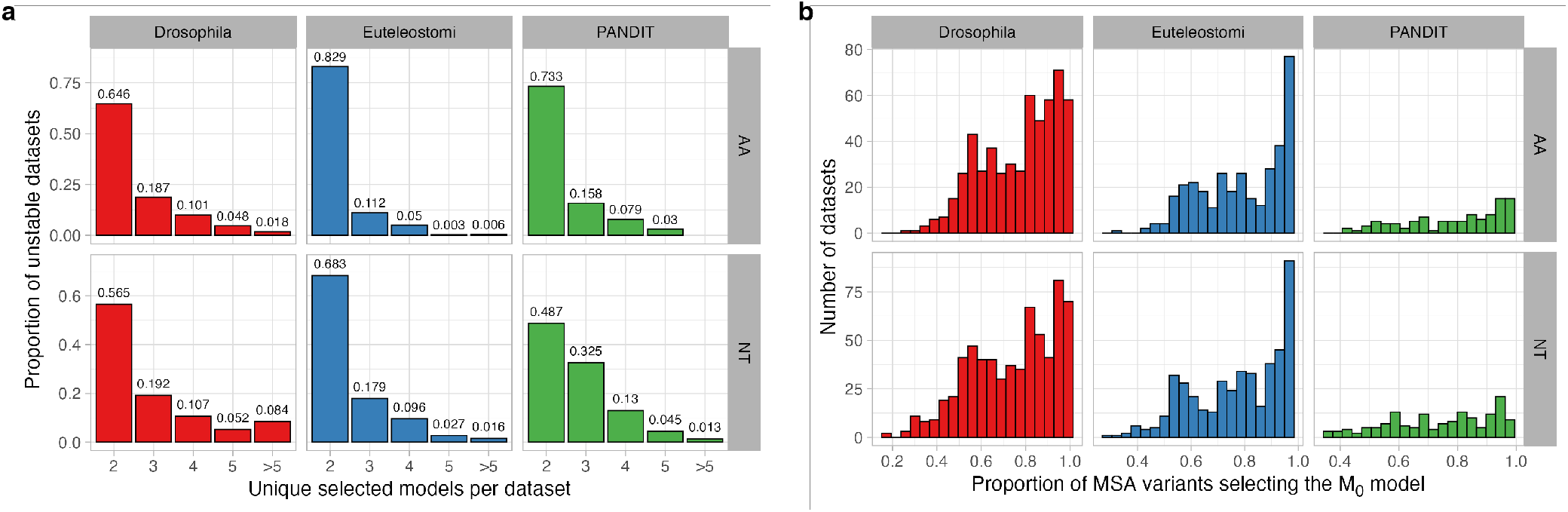
RMS with AIC on unstable datasets. Corresponding figures for model selection with BIC and AICc are in Figure S1). a) Barplot of the total number of unique selected models across the 50 variant MSAs generated for each dataset. Percentages of unstable datasets are shown separately for each dataset source (Drosophila, Euteleostomi, PANDIT) and datatype (AA and NT). b) Histogram of the percentage of MSA variants whose selected model matched *M*_0_ (as determined by the respective information theoretic criterion), the most commonly selected model among all 50 MSA variants (Table 1).

We next assessed, for each unstable dataset, the percentage of MSA variants whose selected model matched the dataset’s most frequently identified best-fitting model, *M*_0_ (Table 1). We observed substantial variation across datasets in how commonly the *M*_0_ model was selected (Figure 2B for AIC and FigureS1b,d for BIC and AICc, respectively). Taken together, these results suggest that there is strong potential for MSA uncertainty to affect model selection, and there is no guarantee that model selection will consistently identify the same best-fitting model for different MSA versions, amino-acid or nucleotide alike, derived from the same underlying data.

We next looked more closely at the differences among selected models for unstable datasets. Specifically, assuming the same datatype (amino-acid or nucleotide), evolutionary models can differ in two primary ways. First, models may have entirely different *Q* matrices, reflecting fundamentally different evolutionary patterns (e.g. `WAG` (Whelan and Goldman 2001) vs. `JTT` (Jones et al. 1992) for amino-acid data). Alternatively, models can have the same underlying *Q* matrix, reflecting a shared evolutionary process, but differ in their additional parameterizations including the specification of stationary frequencies (`F`), presence of among-site rate variation (ASRV, represented here by a four-category discrete Gamma distribution and denoted `+G`), or proportion of invariant sites (`+I`). For example, the two models `WAG+G` and `WAG+G+I` would have the same *Q* matrix but different additional parameterizations.

We therefore asked, specifically among unstable datasets, whether best-fitting models across MSA variants shared the same *Q* matrix (Figure 3 for AIC and Figure S2 for BIC and AICc). To this end, we compared the *Q* matrix selected for each variant MSA to the respective *M*_0_ model’s *Q* matrix, identifying each as either the same or different *Q* matrix.

**Figure 3:**
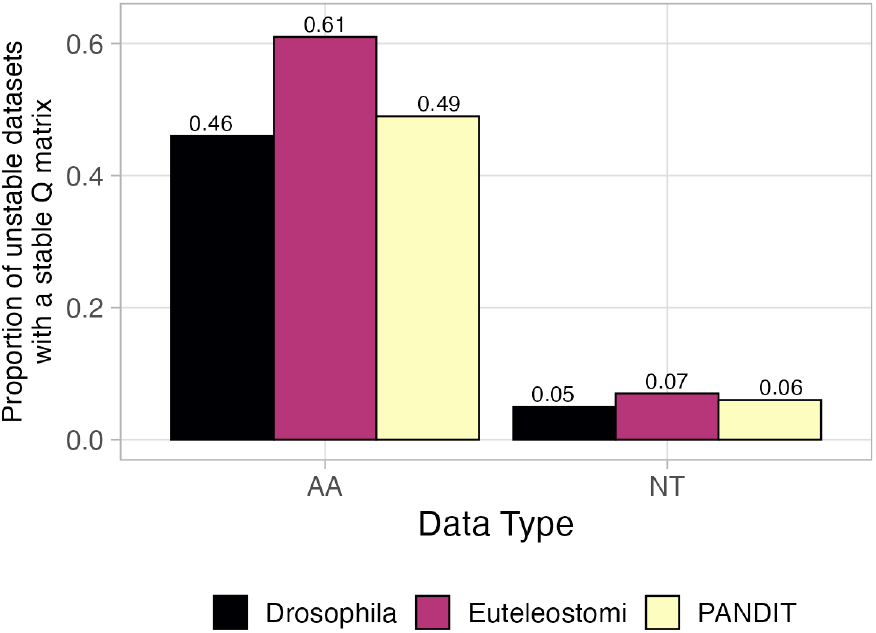
Consistency of *Q* matrices of selected models across MSA variants. “Different *Q* matrices” implies the selected model’s matrix differs from the respective *M*_0_ model’s *Q* matrix, and “Same *Q* matrices” implies the selected model’s matrix matches the respective *M*_0_ model’s *Q* matrix.

As with previous analyses, we observed distinct trends between amino-acid and nucleotide datasets, but overall similar trends among information theoretic criteria and dataset sources examined. For Drosophila and PANDIT amino-acid datasets, roughly half of the selected models shared the *M*_0_ model’s *Q* matrix, but for over 60% of Euteleostomi amino-acid datasets, the selected model’s *Q* matrix matched the *M*_0_ matrix (Figure 3). By contrast, for unstable nucleotide datasets from all data sources, the vast majority of selected *Q* matrices differed from the *M*_0_ model’s *Q* matrix.

One reason why uncertainty in nucleotide MSAs may show increased model selection instability may be the relative model selection procedure itself. The `ModelFinder` algorithm used here for model selection (Kalyaanamoorthy et al. 2017) 88 candidate model parameterizations for nucleotide MSAs, and 168 model parameterizations for amino acid MSAs. More specifically, nucleotide model selection evaluates 21 different *Q* matrices each with four different combinations of `+F` (use observed stationary frequencies), `+I` (proportion of invariant sites), and `+G` (four-category discrete gamma distribution to model among-site rate variation, ASRV) parameterizations, and amino-acid model selection evaluates 22 different *Q* matrices each with six different combinations of `+F`, `+I`, and `+G` parameterizations. As such, there are more model parameterizations per *Q* matrix being evaluated for amino-acid model selection compared to nucleotide model selection. It is therefore possible that this analysis is somewhat biased in favor of amino-acids datasets showing increased *Q* matrix consistency compared to nucleotide datasets simply because there are more models per *Q* matrix. In addition, exchangeability parameters in amino-acid models are not optimized by maximum likelihood, but rather are fixed to *a priori* determined values. By contrast, nucleotide model parameters are all optimized by maximum likelihood and none are *a priori* fixed. This distinction in the source of model parameters may also explain the differences observed between nucleotide and amino-acid datasets.

### Unstable datasets are challenging to distinguish from stable datasets

We next asked whether there are any overall differences between stable and unstable datasets. To this end, we constructed logistic regressions to explore whether MSA stability could be explained by underlying properties of each dataset. We built models for each combination of information theoretic criterion and for each datatype, each considering these four predictors to capture dataset-level information: 1) number of MSA sequences, 2) the mean number of sites among the given dataset’s de-gapped sequences, 3) the mean edit (Hamming) distance between all pairwise comparisons of dataset sequences, and 4) the mean GUIDANCE residue pair scores calculated based on each dataset’s reference MSA (Penn et al. 2010). Smaller mean edit scores indicate overall more similar sequences, and higher mean edit scores indicate more diverged sequences. Similarly, GUIDANCE scores which range between [0, 1] reflect dataset robustness to MSA uncertainty as a whole. Smaller mean GUIDANCE scores indicate that the dataset is challenging to align with high expected uncertainty, and higher mean GUIDANCE scores indicate that the dataset can be more reliably aligned with less expected uncertainty. We calculated edit distances from all-to-all pairwise alignments [constructed with default settings in `MAFFT v7` Katoh and Standley (2013)], and we averaged resulting edit distances to derive a single mean edit distance score for each dataset. GUIDANCE residue pair scores were calculated for the reference MSA against all 49 associated perturbed MSAs, and we averaged all residue pair scores to derive a single mean GUIDANCE score for each dataset.

For all logistic regressions, the mean number of sites and mean number of sequences were significant (all *P* < 10^−5^) predictors of dataset stability but with effect sizes barely different from zero (Figure 4a). By contrast, higher mean GUIDANCE scores were consistently associated with increased log-odds of dataset stability, demonstrating that datasets whose variant MSAs are more similar to another are more likely to be stable, as expected. Surprisingly, the mean edit distance predictor showed opposite trends between amino-acid and nucleotide datasets. For amino-acid data, higher mean edit distance was associated with higher log-odds of dataset stability, except for BIC model selection where this predictor was not significant. For nucleotide data, on the other hand, lower mean edit distances were associated with increased log-odds of dataset stability. Regardless, consistent across information theoretic criteria and datatypes, the mean GUIDANCE score had the largest absolute effect on explaining dataset stability. ROC (receiver operating characteristic) analyses for these logistic regressions revealed an AUC (area under the curve) of at most 0.71 (Figure 4). While better than random chance (AUC=0.5), AUC values ranging from 0.68–0.71 do not suggest a high predictive ability for assessing whether model selection will be sensitive to MSA uncertainty for any given dataset in a real-world analysis. That said, that the predictive ability of these logistic regressions, and similarly AUC scores, may be influenced by having only *N* = 50 perturbed MSAs for each dataset, and exploring more variant MSAs may influence these overall findings. We further note that large ranges of predictor variables (Figure 4b) may also have an impact on resulting AUC values.

**Figure 4:**
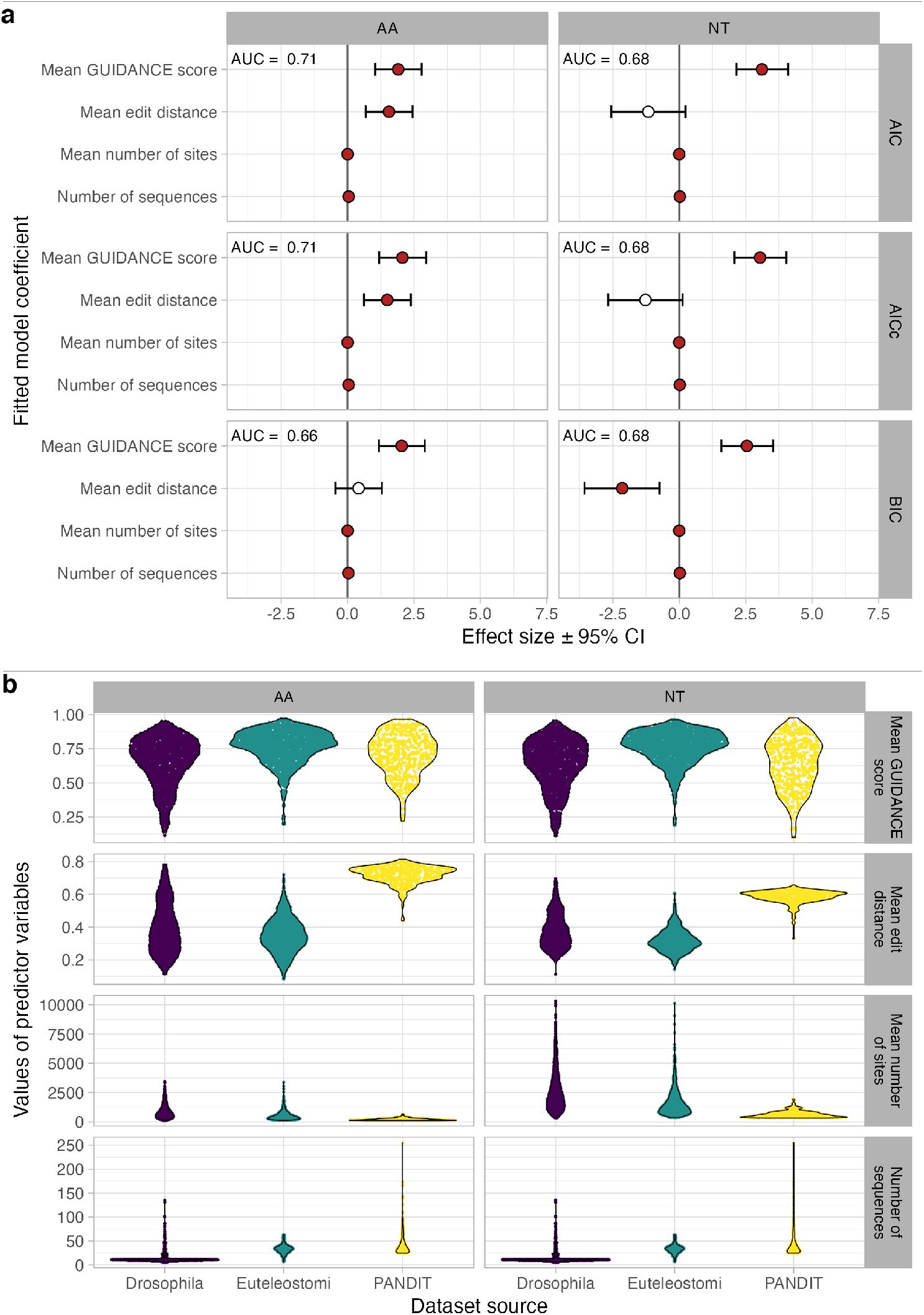
Logistic regression coefficients to explain dataset stability. a) Filled circles represent logistic model coefficients that are significant at *P* ≤ 0.01, and error bars represent the 99% confidence intervals. Values in the top left of each panel represent the AUC from an associated ROC (receiver operating characteristic) analysis. Note that effect sizes for “mean number of sites” and “number of sequences” are significant but extremely close to zero. Dataset sizes in each logistic regressions are equivalent to the number of datasets explored: 236 PANDIT datasets, 1000 Drosophila datasets, and 1000 Euteleostomi datasets, for each of AA and NT datatypes. b) Distributions of predictor variables used in each logistic regression.

We next examined MSA-level properties to ascertain whether more similar variant MSAs were more likely to have the same best-fitting model, and conversely whether more dissimilar variant MSAs are more likely to have different best-fitting models. For this analysis, we considered two common MSA scores, total columns (TC) and sum-of-pairs (SP) scores, to quantify similarity between each dataset’s perturbed MSA and the given reference MSA, using a procedure outlined in Figure 5. For all stable datasets, all perturbed MSA selected models are by-definition the same as the reference MSA’s model (`M`_ref_; Table 1). For each stable dataset, we calculated the SP and TC score between each of the 49 perturbed MSA and the reference MSA (49 total comparisons), finally averaging these scores to derive an overall mean SP and TC score for the dataset. For all unstable datasets, we classified each perturbed MSA either as having the same or different selected model as `M`_ref_, resulting in two groups of MSAs: *differs* `M`_ref_ and *matches* `M`_ref_. We calculated mean TC and SP scores for each of the two groups. Any unstable dataset with fewer than three perturbed MSAs per group was fully excluded from this analysis. In total, these calculations led to three distributions of mean dataset SP and TC scores for each of the *stable, differs*, and *matches* groups (Figure 6 for AIC and Figure S3a,b for BIC and AICc, respectively).

**Figure 5:**
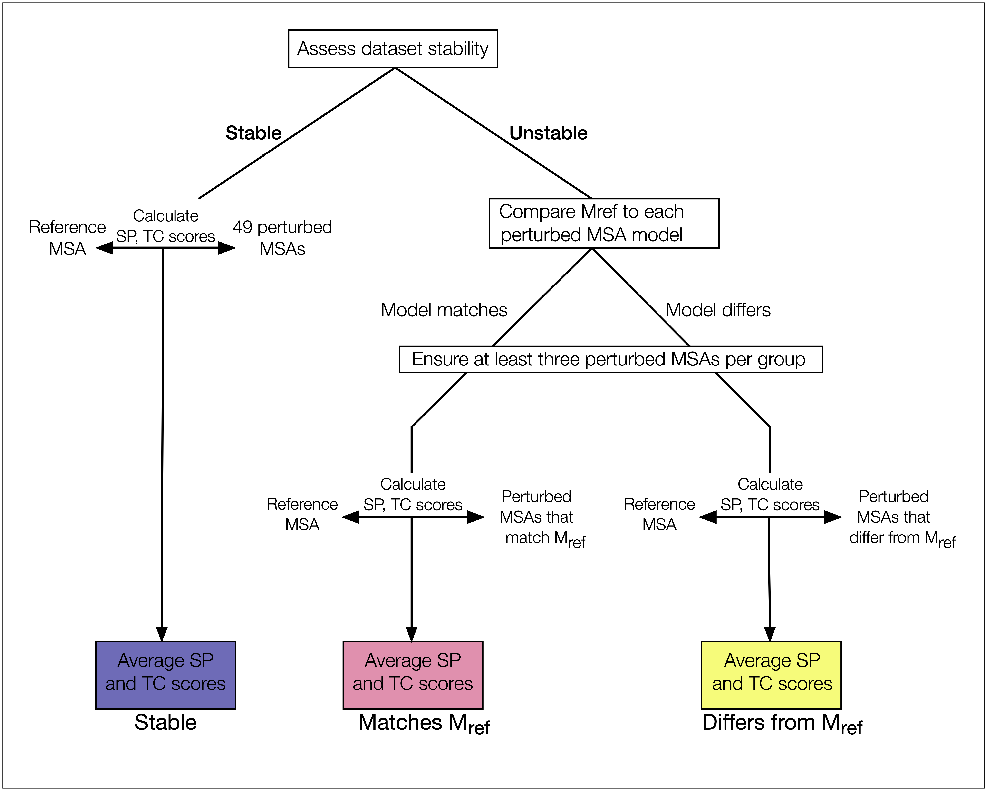
Flowchart showing procedure for grouping MSAs to analyze SP and TC scores. Endpoints in the flowchart are colored according to the same groupings shown in Figure 6.

**Figure 6:**
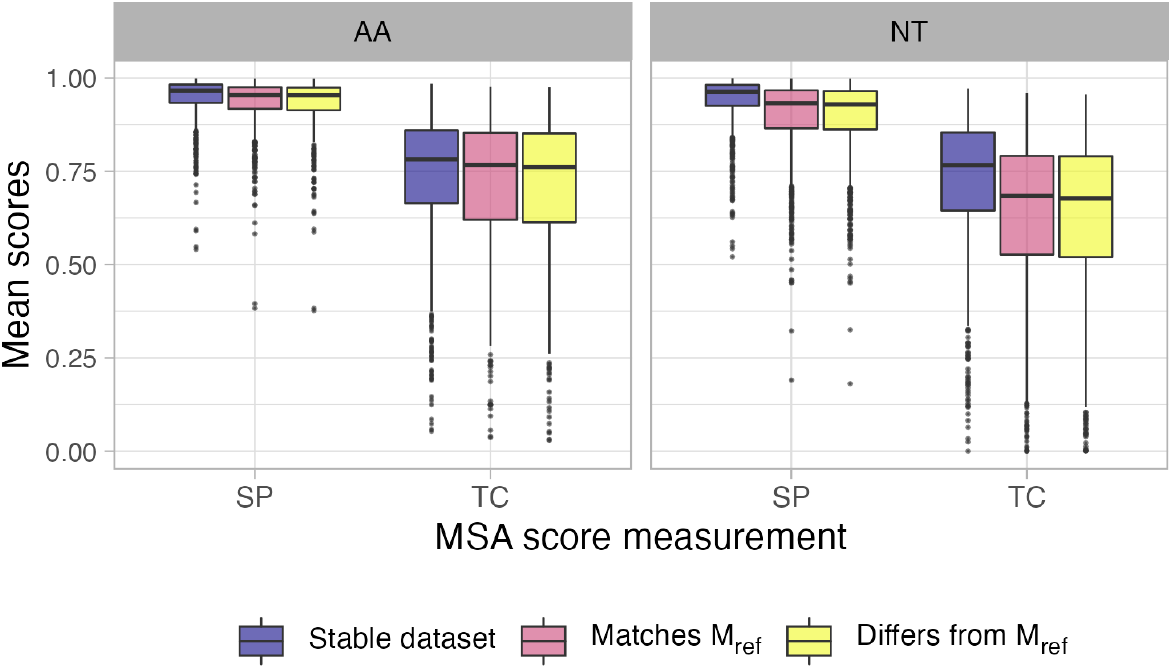
Mean SP and TC scores across variant MSA groups for model selection with AIC. Values in each boxplot represent mean dataset SP and TC scores across MSA variant groupings. Corresponding figures for model selection with BIC and AICc are in Figure S3.

For each datatype (NT and AA) and MSA score (SP and TC), we built an ANOVA to assess the mean score differences among the three groupings, applying a post-hoc Tukey test to compare differences among groupings (Table 2 for AIC, and Tables S1-2 for BIC and AICc, respectively). Overall, we observed no significant differences the *differs* and *matches* groupings for all information theoretic criteria (*differs-matches* comparisons in Table 2 and Tables S1-2). By contrast, all *stable-differs* and all except one *stable-matches* comparison were significant at *P* ≤ 0.01, with one exception: The *stable-matches* comparison of TC scores on amino-acid datasets with AIC model selection was significant only at the level *P* ≤ 0.05 (Table 2). Moreover, all effect sizes were fairly small, except for significant nucleotide TC score comparisons. In total, these results support findings shown in Figure 4 that MSAs are overall more similar for stable datasets compared to unstable datasets, but specifically predicting whether two given variant MSAs will select the same evolutionary model may not be possible.

**Table 2:** SP and TC score comparisons among variant MSA groups for model selection with AIC. The “Comparison” column indicates the difference in effect sizes between the given groupings, which were compared in a post-hoc Tukey test with corrected P-values. Rows with adjusted *P* ≤ 0.01 are shown in bold. Corresponding results for model selection with BIC and AICc are in Tables S1 and S2, respectively.

| Datatype | Score type | Comparison | Estimate (99% CI) |
| --- | --- | --- | --- |
| AA | SP | differs-matches | -0.002 (-0.011, 0.007) |
| **AA** | **SP** | **stable-matches** | **0.014 (0.006, 0.022)** |
| **AA** | **SP** | **stable-differs** | **0.016 (0.008, 0.024)** |
| AA | TC | differs-matches | -0.007 (-0.033, 0.018) |
| AA | TC | stable-matches | 0.022 (-0.001, 0.045) |
| **AA** | **TC** | **stable-differs** | **0.029 (0.006, 0.052)** |
| NT | SP | differs-matches | -0.003 (-0.015, 0.01) |
| **NT** | **SP** | **stable-matches** | **0.043 (0.031, 0.055)** |
| **NT** | **SP** | **stable-differs** | **0.046 (0.034, 0.058)** |
| NT | TC | differs-matches | -0.006 (-0.032, 0.021) |
| **NT** | **TC** | **stable-matches** | **0.093 (0.066, 0.119)** |
| **NT** | **TC** | **stable-differs** | **0.098 (0.071, 0.125)** |

## Discussion and Conclusions

We have conducted a large-scale empirical data analysis to ascertain the extent to which the results of relative model selection may be influenced by MSA uncertainty. We assessed the consistency of model selection, for three different commonly-used information theoretic criteria (AIC, BIC, AICc), across variant MSAs created from the same dataset, finding that model selection is often sensitive to MSA uncertainty. As has been often observed in previous studies, we observed very few practical differences among results from different information theoretic criteria used in model selection (Ripplinger and Sullivan 2008; Abadi et al. 2019). Furthermore, we have shown that it is challenging, if at all possible, to predict the specific circumstances under which MSA uncertainty influences model selection, although we do observe that model selection is, as expected, more consistent when the MSA itself is more reliable (Figure 4). In total, these results build on previous findings that MSA uncertainty has the potential to influence even more stages of phylogenetic analyses than has been previously recognized.

We note that one limitation of this study is that we only explore the influence of MSA uncertainty on relative model selection and not other approaches to identifying best-fitting models, including tests of model adequacy (Goldman 1993a,b; Duchêne et al. 2018), Bayesian assessments of absolute model fit (Brown 2014; Lewis et al. 2014), or more recently-developed machine-learning methods for model selection (Abadi et al. 2020). Indeed, it may be possible that other approaches to model selection are more robust to MSA uncertainty, but the computational demands of these methods prohibit similar large-scale benchmarking. As the computational efficiency of these methods improves, it will be possible to thoroughly examine the potential effects of MSA uncertainty on model adequacy approaches.

How can the effects of MSA uncertainty be mitigated for relative model selection? In other scenarios in phylogenetic methods where MSA uncertainty is a known confounding factor, researchers will sometimes filter and/or mask MSAs to remove columns and/or residues that are suspected to have been poorly aligned. However, there have been mixed results on whether such actions will improve analyses. For example, in the context of positive selection, Privman et al. (2012) found that masking poorly aligned residues may reduce false positives, but Spielman et al. (2014) found that there was limited practical utility to masking residues. Similarly, in the context of phylogenetic reconstruction, Sela et al. (2015) found that MSA filtering may improve inferences, but Tan et al. (2015) found that MSA filtering more frequently worsens inferences. Overall, these studies point to the conclusion that MSA filtering can reduce spurious inferences, but may also reduce power to the detriment of overall performance. In spite of the lackluster promise shown by other applications of MSA filtering, examining the effect of MSA filtering on model selection itself may begin to reveal the source of its uncertainty. Another potential option for mitigating the effects of MSA uncertainty more generally may be to pursue a model averaging and/or mixture modeling strategy (Posada and Buckley 2004; Si Quang et al. 2008; Bouckaert and Drummond 2017), for example by jointly considering candidate models identified across variant MSAs, during phylogenetic reconstruction. Alternatively, it may be possible to address MSA uncertainty by averaging across a candidate set of MSAs, which has been previously shown to improve accuracy in phylogenetic reconstruction (Sela et al. 2015), and selecting a model for subsequent analyses based on the averaged MSA rather than from a single variant MSA.

While it may be possible to ameliorate the influence of MSA uncertainty on relative model selection, we must also ask: Do we need to mitigate this issue in the first place? For example, recent studies have shown that, for both nucleotide and amino-acid models, the model selection procedure itself may not be a critical step in phylogenetic reconstruction, since different models with extreme differences in relative fit may not actually result in systematically different results (Spielman and Kosakovsky Pond 2018; Abadi et al. 2019; Spielman 2020) although how the precise model used may influence branch length and/or divergence estimation remains an important question (Abadi et al. 2019, 2020). As such, if distinct models may yield highly similar inferences, optimizing the model selection procedure itself has diminishing returns. If models themselves are overly similar, there may be little practical consequence to the observed uncertainty in model selection.

Considering the results of this study in the context of literature questioning the utility of model selection itself, we suggest that, rather expending additional efforts to perfect existing paradigms in phylogentic modeling, the time has come to explore novel modeling paradigms. For example, one reason that all models may yield highly similar inferences is that, while they may have somewhat different focal parameters, virtually all commonly-used models, and certainly all those evaluated by model selection, adhere to the same underlying mechanism: A stationary, time-reversible, and homogeneous Markov process that assumes sites evolve independently (e.g. no epistasis) (Naser-Khdour et al. 2019). We conclude that concerted efforts to develop and extend these paradigms while facilitating widespread familiarity represent the most promising next steps in advancing phylogenetic methods.

## Methods

### Construction of perturbed alignments

We analyzed multiple sequence alignments (MSAs) from two databases, Selectome v6 (Moretti et al. 2014) and PANDIT (Whelan 2006), specifically considering only those MSAs which had fully compatible amino-acid and nucleotide versions. This approach allowed us to use the same underlying biological data to examine the robustness of relative model selection when performed on both nucleotide and amino-acid MSAs. From Selectome, we analyzed 1000 randomly selected Drosophila alignments and 1000 randomly selected Eutelestomi MSAs, ensuring that each MSA corresponded to a different gene (i.e., only a single transcript for a given gene was analyzed). From PANDIT, we analyzed all MSAs which contained at least 25 sequences and whose shortest protein sequence contained at least 100 amino-acids, which totalled 254 PANDIT MSAs.

After collecting these datasets, we removed all gaps from these MSAs. We refer to each resulting set of unaligned orthologs as a “dataset.” For each nucleotide and amino-acid version of each dataset, we used a `GUIDANCE2`-based approach to generate 50 distinct MSAs (Sela et al. 2015; Spielman et al. 2014). We created perturbed alignments by varying the guide tree and gap penalty parameters provided to the `MAFFT v7.407` aligner (Katoh and Standley 2013). In this procedure, a “reference” MSA is first made using default parameters in `MAFFT`. Bootstrapped versions of this reference MSA are used to create guide trees for the progressive alignment with `FastTree2` (Price et al. 2010). Each unique guide tree is then supplied to `MAFFT` as a guide tree to construct a perturbed MSA from the underlying data. As in `GUIDANCE2`, the `MAFFT` gap opening penalty *X* for specified each perturbed MSA was randomly drawn from the uniform distribution *X* ∼ *U* (1, 3). We created 49 perturbed MSAs for each dataset, leading to a total of 50 distinct MSAs (including the reference MSA) for each dataset. We ensured that all MSAs for a given dataset were fully unique.

### Relative model selection

We used `ModelFinder` (Nguyen et al. 2015; Kalyaanamoorthy et al. 2017) to conduct relative model selection, using the `-m TEST` flag to perform a standard model selection similar to `jModelTest` and `ProtTest` (Darriba et al. 2012, 2011). This procedure evaluates 88 candidate model parameterizations for nucleotide MSAs, and 168 model parameterizations for amino acid MSAs. More specifically, nucleotide model selection evaluates 21 different *Q* matrices each with four different combinations of `+F` (use observed stationary frequencies), `+I` (proportion of invariant sites), and `+G` (four-category discrete gamma distribution to model among-site rate variation, ASRV) parameterizations, and amino-acid model selection evaluates 22 different *Q* matrices each with six different combinations of `+F`, `+I`, and `+G` parameterizations. For each dataset, we identified the best-fitting model under each of the three information theoretic criterion AIC (Akaike Information Criterion), AICc (small-sample Akaike Information Criterion), and BIC (Bayesian Information Criterion).

## Declarations

### Ethics approval and consent to participate

Not applicable.

### Consent for publication

Not applicable.

### Availability of data and materials

All code used in this study is freely available from https://github.com/spielmanlab/alignment_models. *Upon acceptance of this manuscript, the code repository will be indexed with a permanent DOI in Zenodo, and all data (variant MSAs) will be deposited in an appropriate data repository such as FigShare or Data Dryad*.

### Authors’ contributions

SJS conceived of and designed the study. SJS and MM performed data analysis. SJS wrote the paper. All authors have read and approved the manuscript.

## Acknowledgements

High-performance computing was conducted, in part, using the Rowan University Computing Cluster (RUCC).

