## Supplemental Materials for "Relative model selection of evolutionary substitution models can be sensitive to multiple sequence alignment uncertainty"

### Supplementary Figures

#### Figure S1

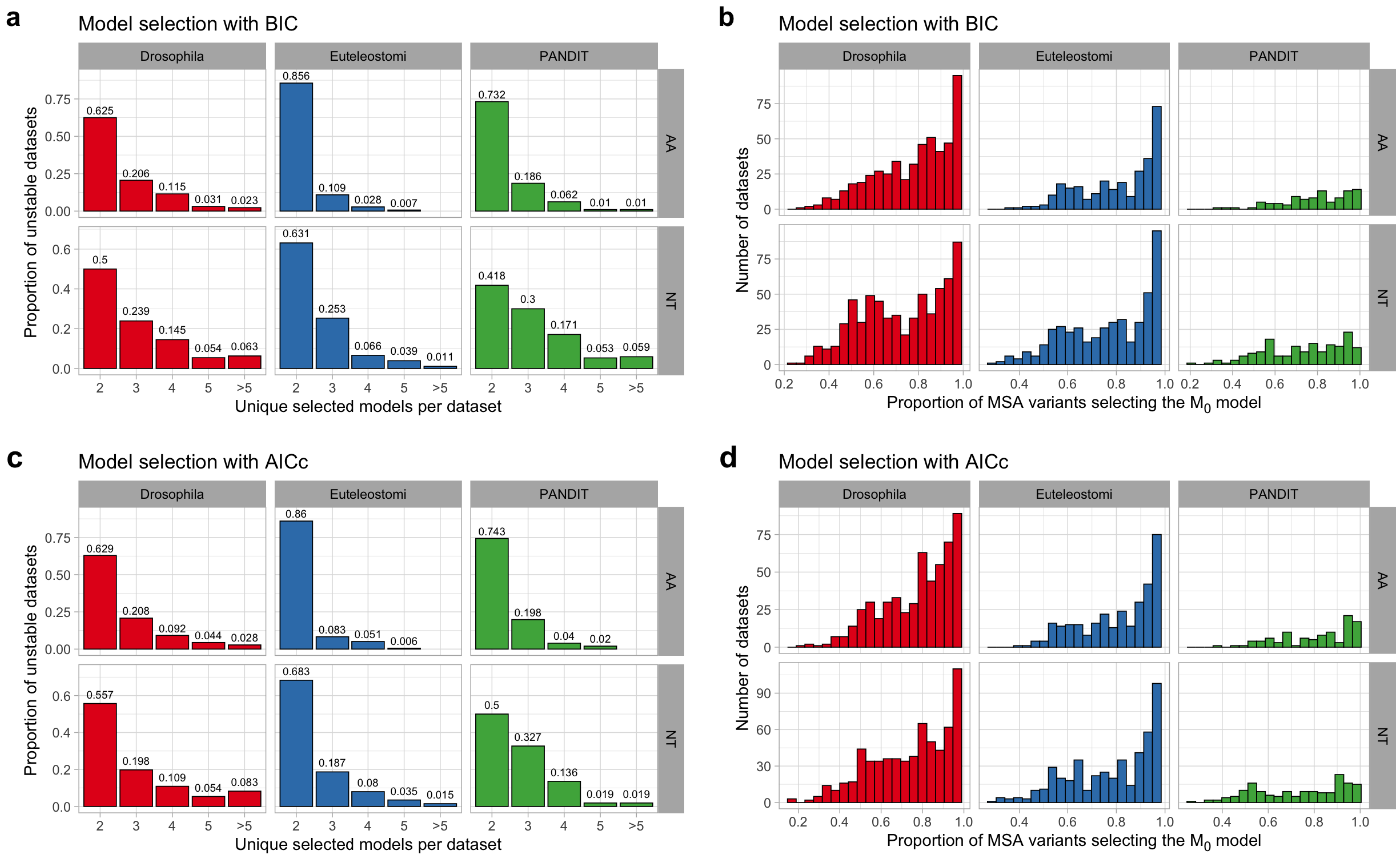

### Figure S2

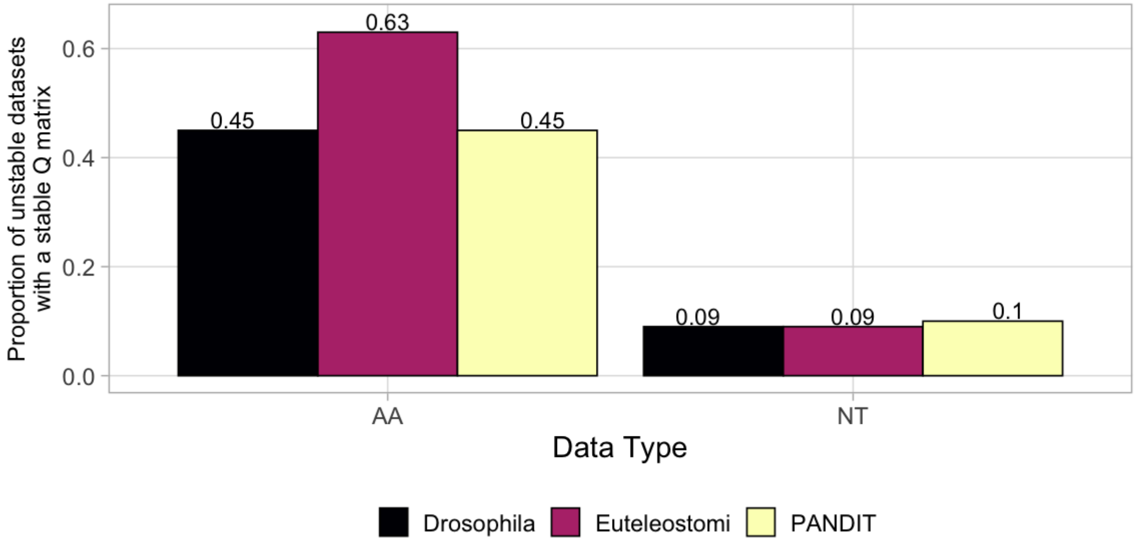


### Figure S3

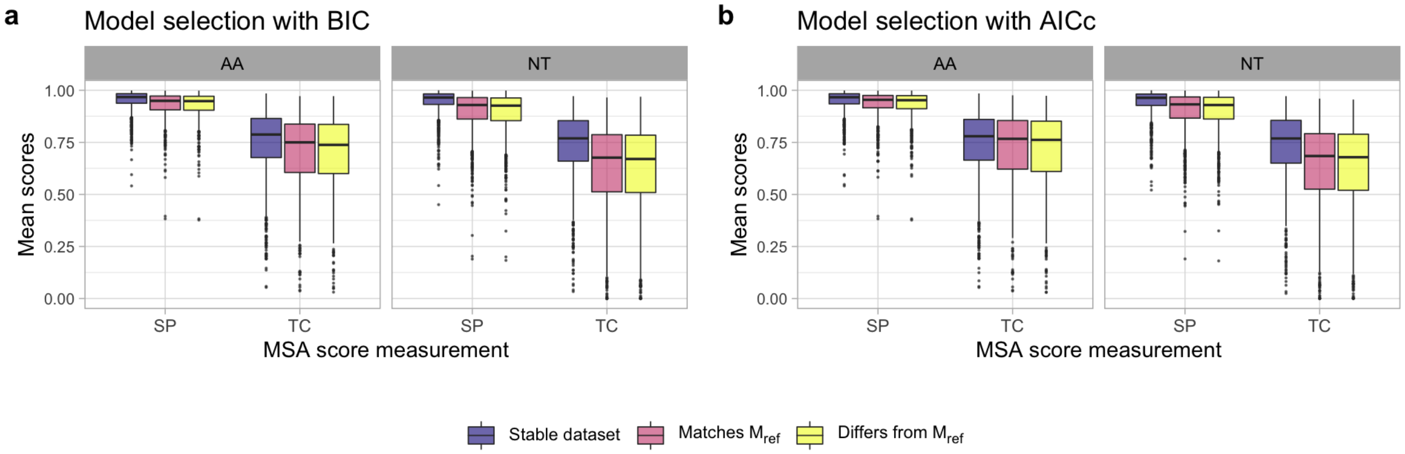

### Supplementary Tables

#### Table S1

SP and TC scores comparisons among groups

Model selection with BIC

| Datatype | Score type | Comparison | Effect size (99% CI) | Adjusted P-value |
|---|---|---|---|---|
| AA | SP | differs-matches | -0.002 (-0.011, 0.008) | 0.867 |
| **AA** | **SP** | **stable-matches** | **0.023 (0.015, 0.032)** | **0.000** |
| **AA** | **SP** | **stable-differs** | **0.025 (0.017, 0.033)** | **0.000** |
| AA | TC | differs-matches | -0.006 (-0.034, 0.021) | 0.769 |
| **AA** | **TC** | **stable-matches** | **0.049 (0.026, 0.073)** | **0.000** |
| **AA** | **TC** | **stable-differs** | **0.056 (0.032, 0.079)** | **0.000** |
| NT | SP | differs-matches | -0.003 (-0.015, 0.009) | 0.739 |
| **NT** | **SP** | **stable-matches** | **0.05 (0.038, 0.062)** | **0.000** |
| **NT** | **SP** | **stable-differs** | **0.053 (0.041, 0.065)** | **0.000** |
| NT | TC | differs-matches | -0.007 (-0.033, 0.019) | 0.699 |
| **NT** | **TC** | **stable-matches** | **0.105 (0.079, 0.132)** | **0.000** |
| **NT** | **TC** | **stable-differs** | **0.112 (0.086, 0.139)** | **0.000** |

### Table S2

SP and TC scores comparisons among groups

Model selection with AICc

| Datatype | Score type | Comparison | Effect size (99% CI) | Adjusted P-value |
|---|---|---|---|---|
| AA | SP | differs-matches | -0.002 (-0.012, 0.007) | 0.762 |
| **AA** | **SP** | **stable-matches** | **0.016 (0.008, 0.025)** | **0.000** |
| **AA** | **SP** | **stable-differs** | **0.019 (0.01, 0.027)** | **0.000** |
| AA | TC | differs-matches | -0.008 (-0.034, 0.018) | 0.653 |
| **AA** | **TC** | **stable-matches** | **0.025 (0.001, 0.048)** | **0.006** |
| **AA** | **TC** | **stable-differs** | **0.033 (0.009, 0.056)** | **0.000** |
| NT | SP | differs-matches | -0.002 (-0.014, 0.01) | 0.831 |
| **NT** | **SP** | **stable-matches** | **0.044 (0.032, 0.056)** | **0.000** |
| **NT** | **SP** | **stable-differs** | **0.046 (0.034, 0.058)** | **0.000** |
| NT | TC | differs-matches | -0.006 (-0.032, 0.02) | 0.788 |
| **NT** | **TC** | **stable-matches** | **0.095 (0.068, 0.121)** | **0.000** |
| **NT** | **TC** | **stable-differs** | **0.101 (0.074, 0.127)** | **0.000** |
